## Supplemental data for "Mitochondrial Aurora kinase A induces mitophagy by interacting with MAP1LC3 and Prohibitin 2"

**Giulia Bertolin et al.**

### Supplementary information

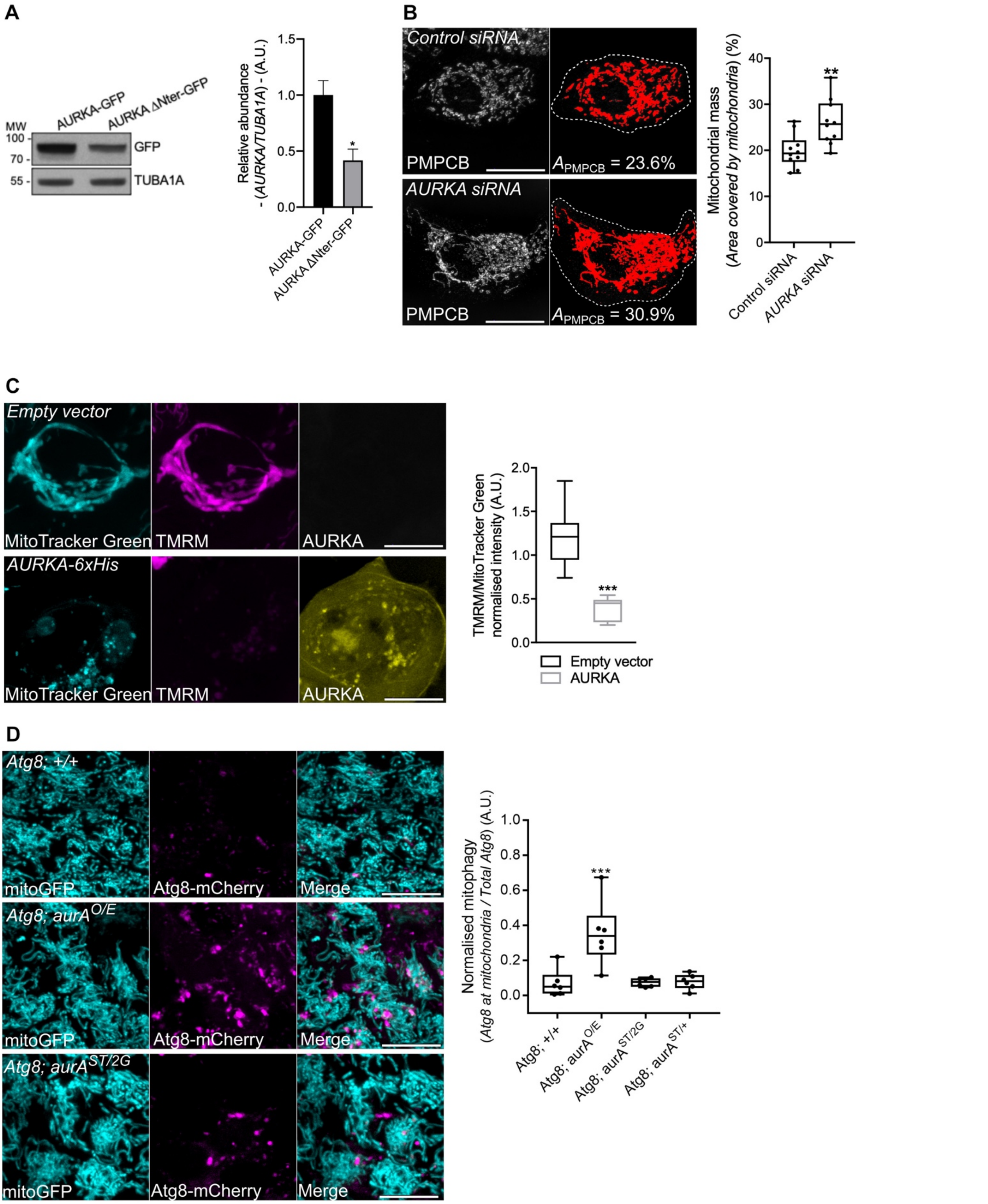


**Supplementary Fig. 1 The over-expression of AURKA induces ΔΨ loss and increases the mitochondrial localisation of Atg8 in *Drosophila melanogaster*.** (A) Representative immunoblot and quantification of the normalised abundance of AURKA-GFP or AURKA ΔNter-GFP in total lysates of HEK293 cells. Loading control: TUBA1A. n=3 independent experiments. (B) Loss of mitochondrial staining (threshold mask and corresponding quantification) of T47D cells transfected with a control or an *AURKA*-specific siRNA. Mitochondria were stained with an anti-PMPCB antibody. $A_{PMPCB}$= PMPCB-labeled mitochondrial area normalised against total cell area (%). *n*=10 cells per condition from one representative experiment (of three). (C) (Left panels) Representative images of MitoTracker Green, TMRM and AURKA-iRFP670 and (right panels) quantification of the TMRM/MitoTracker Green ratio in cells transfected with an empty vector or with AURKA-iRFP670. *n*=10 images per condition on one representative experiment (of three). mitoGFP and MitoTracker Green were pseudocoloured cyan, AURKA-iRFP670 was pseudocoloured yellow, and Atg8-mCherry and TMRM were pseudocoloured magenta. (D) (Left panels) Representative images of a mitochondrially-targeted GFP (mitoGFP) and Atg8-mCherry, and (right panels) corresponding quantification of the number of Atg8-mCherry spots co-localising with mitoGFP and normalised to the total number of Atg8-mCherry spots in a wild-type background (Atg8; +/+), gain-of-function (Atg8; $aurA^{O/E}$), AURKA null ($aurA^{ST/2G}$) and heterozygous ($aurA^{ST/+}$) *Drosophila* mutants. *n*=60 images per condition from six independent pupae obtained from three independent crossings. Scale bar: 10µm. Data extend from the min to max. ***P*<0.01, ****P*<0.001 compared to the 'Control siRNA' (B), 'Empty vector' (C), or the the 'Atg8; +/+' (D) conditions. A.U.: arbitrary units.

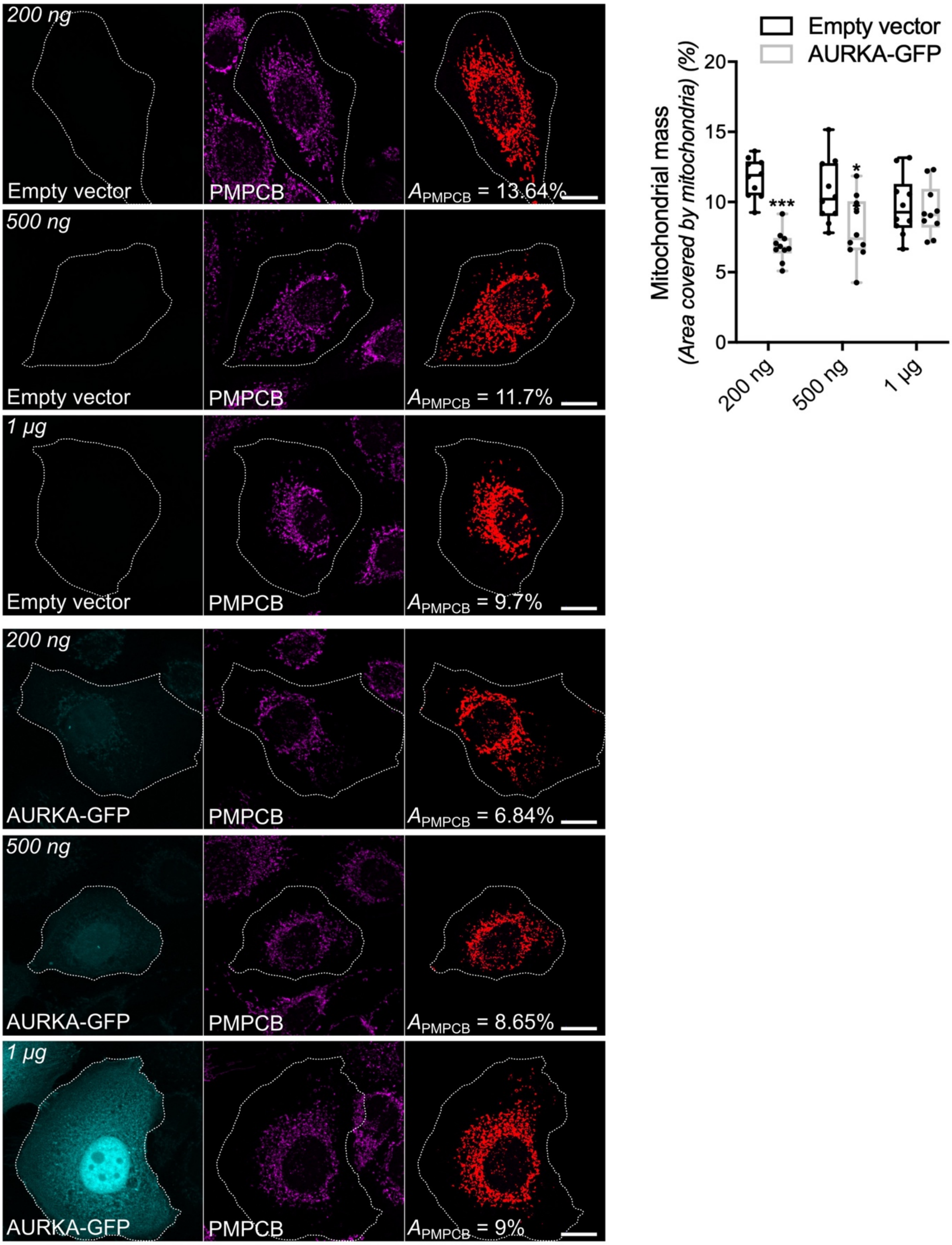


**Supplementary Fig. 2 AURKA-driven mitochondrial mass loss depends on AURKA expression levels.** Representative images and loss of mitochondrial staining (threshold mask and corresponding quantification) of HMLE epithelial cells nucleofected with 200 ng, 500 ng or 1 µg of an empty vector or a plasmid encoding AURKA-GFP. Mitochondria were stained with an anti-PMPCB antibody. The GFP channel was pseudocoloured cyan, PMPCB was pseudocoloured magenta. $A_{PMPCB}$= PMPCB-labeled mitochondrial area normalised against total cell area (%). *n*=10 cells per condition from one representative experiment (of three). Scale bar: 10µm. Data extend from min to max. *$P$<0.05, ***$P$<0.001 compared to each corresponding 'Empty vector' condition.

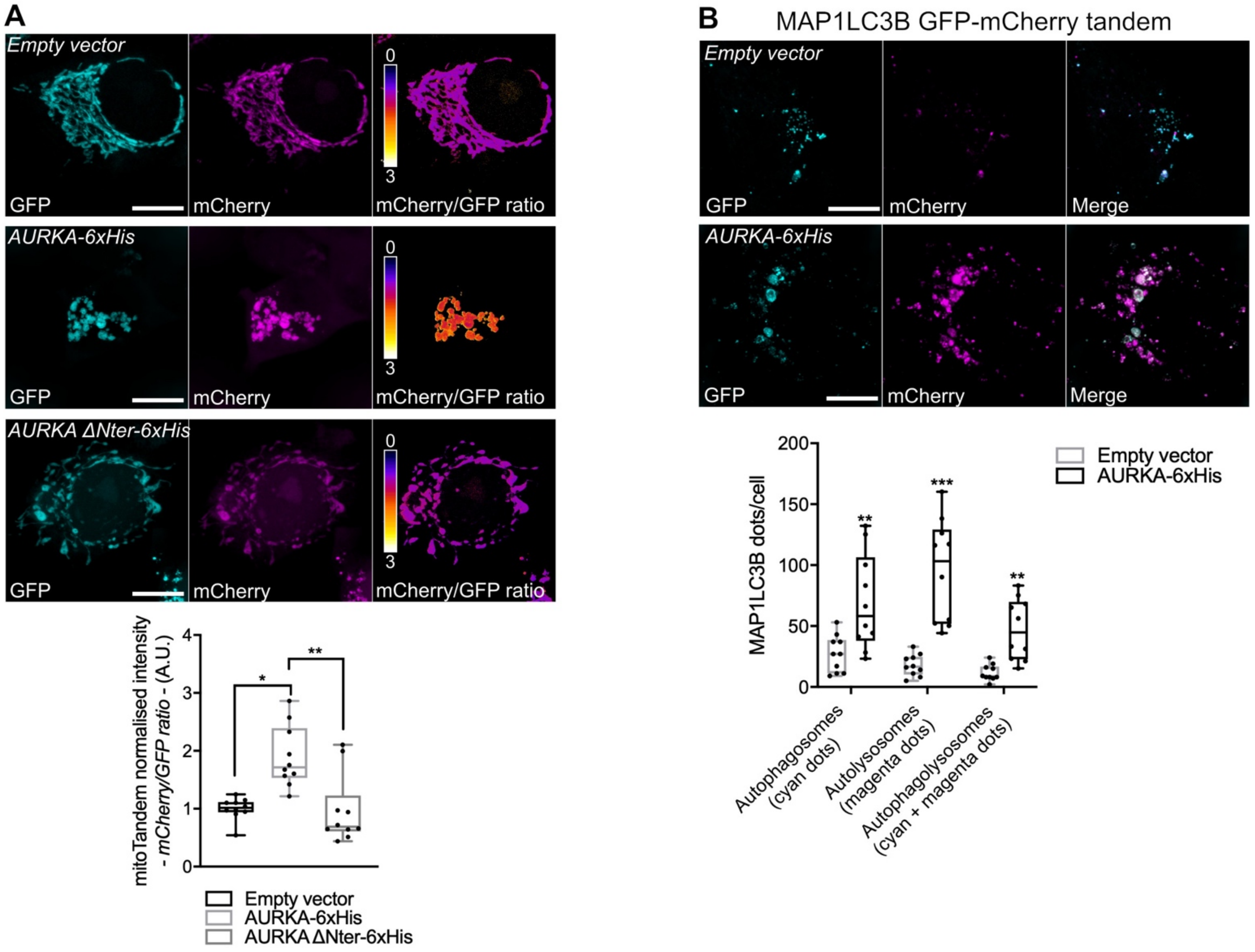


**Supplementary Fig. 3 The over-expression of AURKA induces an autophagy-related drop of mitochondrial pH.** (A) (Upper panels) Representative images of a mitochondrially-targeted GFP/mCherry fluorescent tandem co-transfected with an empty vector, AURKA-6xHis or AURKA ΔNter-6xHis plasmids. Pseudocolour scale: mCherry/GFP ratio. (Lower panel) Quantification of the mCherry/GFP fluorescence intensity ratio. *n*=10 images per condition on one representative experiment (of three). (B) (Upper panels) Representative images of a GFP/mCherry-MAP1LC3 fluorescent tandem co-transfected with an empty vector or a AURKA-6xHis plasmid. (Lower panel) Quantification of the number of MAP1LC3 fluorescent dots in the GFP channel (autophagosomes), in the mCherry one (autolysosomes) or in both fluorescent channels (autophagolysosomes). *n*=10 images per condition on one representative experiment (of three). GFP was pseudocoloured cyan and mCherry was pseudocoloured magenta. Scale bar: 10µm. Data extend from the min to max. **P*<0.05, ***P*<0.01, ****P*<0.001 compared to the 'Empty vector' (A), or each corresponding 'Empty vector' (B) conditions. A.U.: arbitrary units.

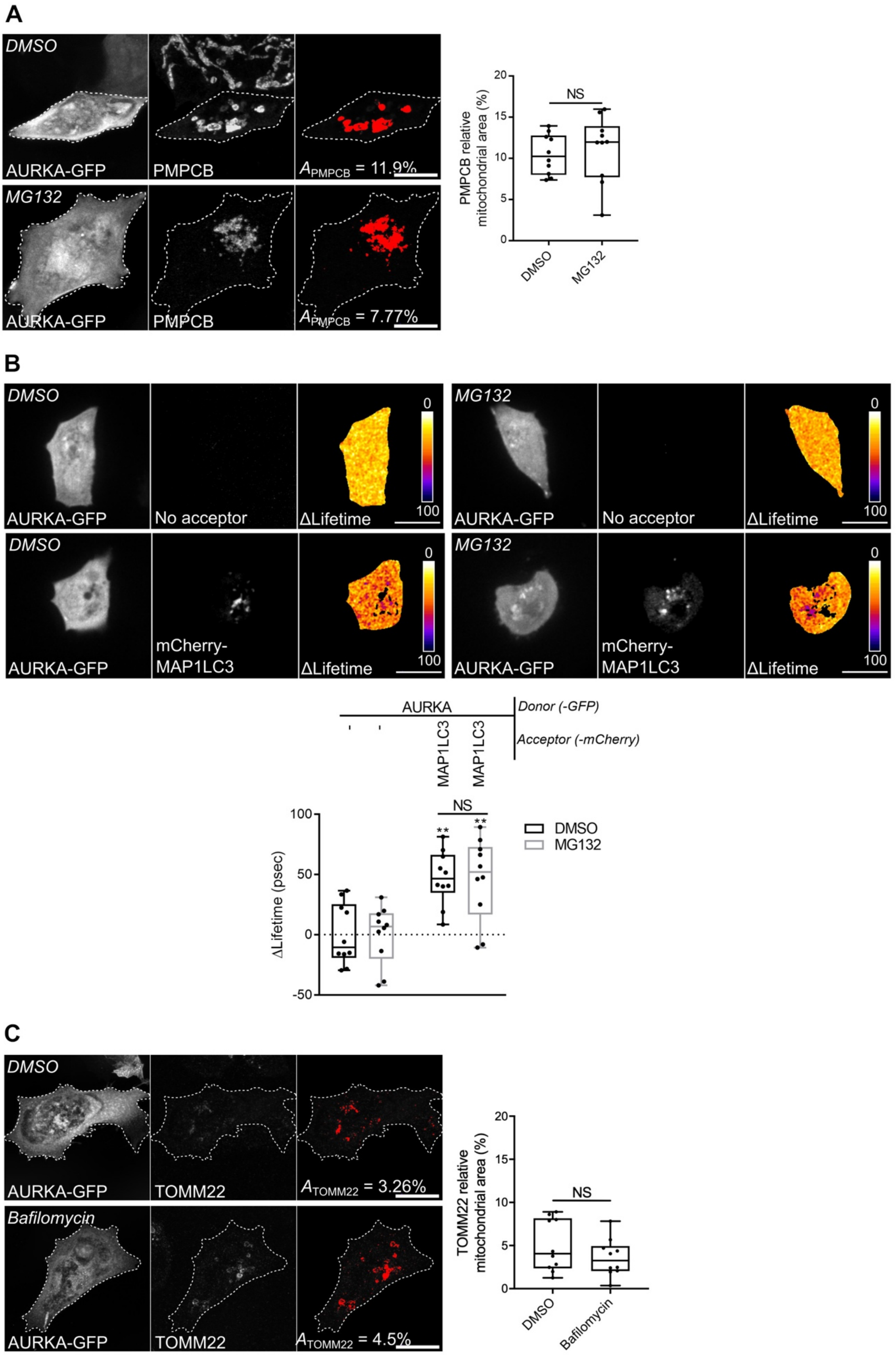


**Supplementary Fig. 4. Proteasomal inhibition does not prevent the loss of matrix markers and the interaction of AURKA with MAP1LC3.** (A) Loss of PMPCB staining (threshold mask and corresponding quantification) in MCF7 cells transfected with AURKA-GFP and treated with MG132. (B) Representative FRET by FLIM images and corresponding analyses on MCF7 cells expressing AURKA-GFP alone or together with mCherry-MAP1LC3B, and treated with DMSO or MG132. Pseudocolour scale: pixel-by-pixel ΔLifetime. (C) Loss of TOMM22 staining (threshold mask and corresponding quantification) in MCF7 cells transfected with AURKA-GFP and treated with DMSO or Bafilomycin. *A*=mitochondrial area normalised against total cell area (%). *n*=10 cells per condition from one representative experiment (of three). Data extend from the min to the max. Scale bar: 10µm. ***P* < 0.01 against each corresponding 'AURKA-No acceptor' condition. NS: not significant.

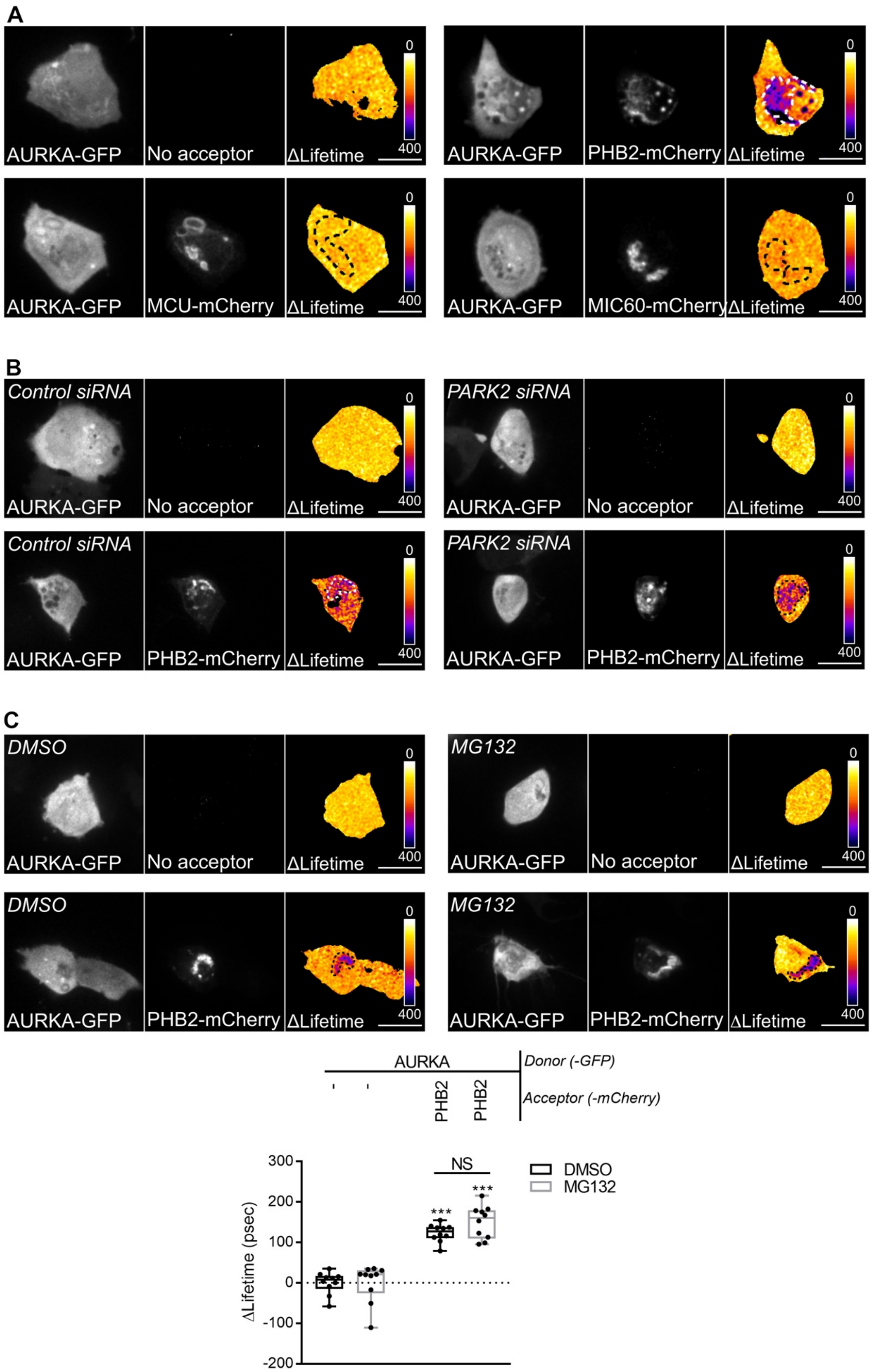


**Supplementary Fig. 5. The AURKA-PHB2 interaction is specific, and is maintained upon *PARK2* silencing and proteasomal inhibition.** (A) Representative FRET by FLIM images of MCF7 cells expressing AURKA-GFP alone or together with PHB2-mCherry, MCU-mCherry or MIC60-mCherry. (B) Representative FRET by FLIM images of MCF7 cells expressing AURKA-GFP alone or together with PHB2-mCherry, and co-transfected with a control- or a *PARK2*-specific siRNA. (C) FRET by FLIM analyses of MCF7 cells expressing AURKA-GFP alone or together with PHB2-mCherry, and treated with DMSO or with MG132. $n$=10 cells per condition from one representative experiment (of three). Data extend from the min to the max. Pseudocolour scale: pixel-by-pixel ΔLifetime. Scale bar: 10µm. ***$P$<0.001 compared to each corresponding 'No acceptor' condition.

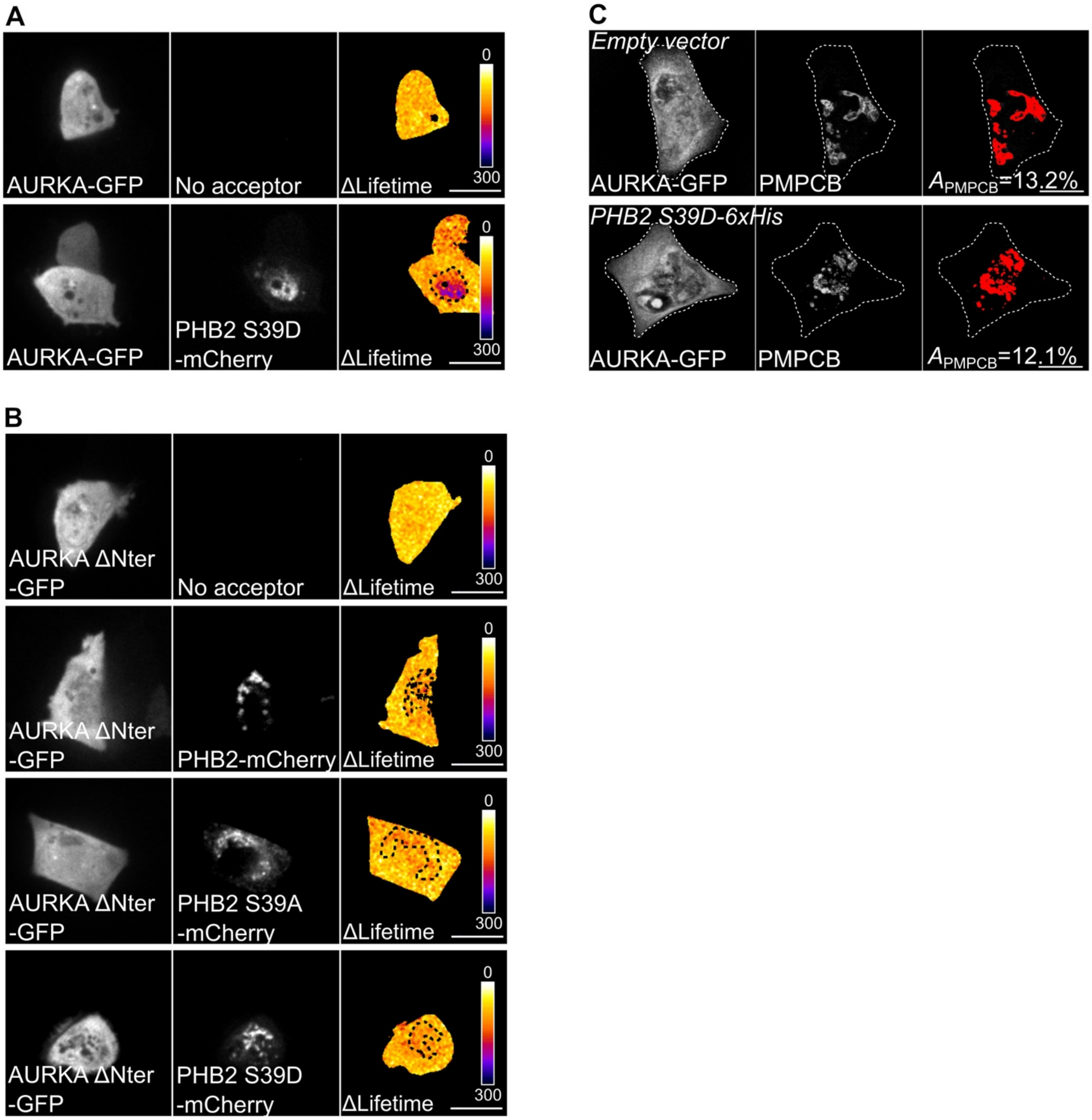


**Supplementary Fig. 6. The AURKA-PHB2 interaction is maintained with PHB2 S39D, but not with AURKA ΔNter.** (A-B) Representative FRET by FLIM images of MCF7 cells expressing AURKA-GFP alone or together with PHB2 S39D-mCherry (A), or expressing AURKA ΔNter alone or with PHB2-mCherry, PHB2 S39A-mCherry or PHB2 S39D-mCherry (B). Pseudocolour scale: pixel-by-pixel ΔLifetime. (C) Loss of PMPCB staining (threshold mask) in MCF7 cells co-transfected with AURKA-GFP and an empty vector or PHB2 S39D-6xHis, as indicated. *A*=mitochondrial area normalised against total cell area (%). Scale bar: 10µm.

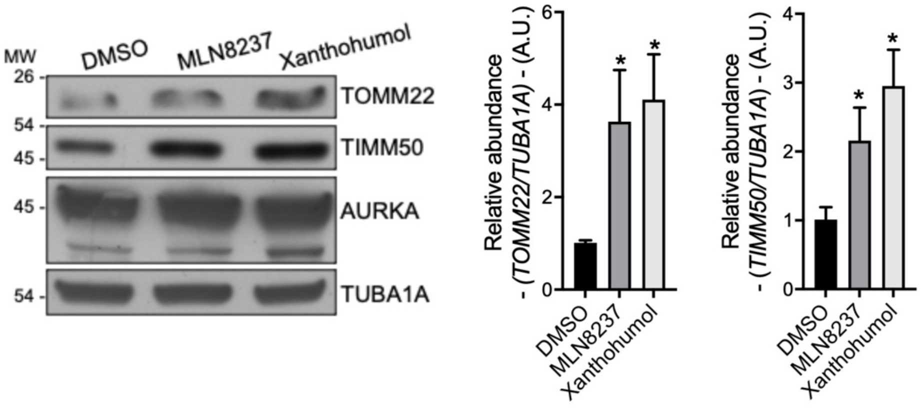


**Supplementary Fig. 7. Xanthohumol and MLN8237 impair mitophagy in AURKA-overexpressing cells.** Representative immunoblot and quantification of the normalised abundance of TOMM22 and TIMM50 in total lysates of HEK293 cells overexpressing AURKA-6xHis and treated with DMSO, MLN8237 or Xanthohumol for 24h. Loading control: TUBA1A. *n*=3 independent experiments. Data represent means ± s.e.m. **P* < 0.05, ***P* < 0.01, ****P*<0.001 compared to each corresponding 'DMSO' condition. A.U.: arbitrary units.

#### Fig. 1A

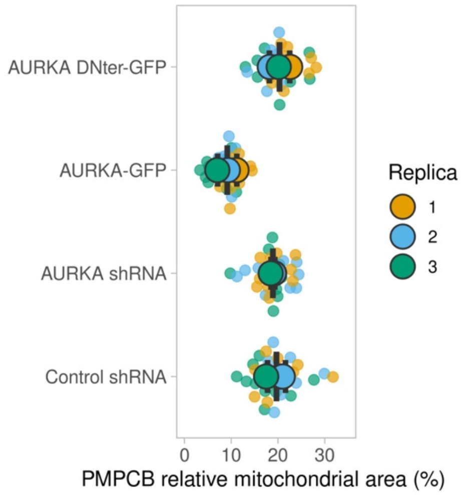


#### Fig. 1E

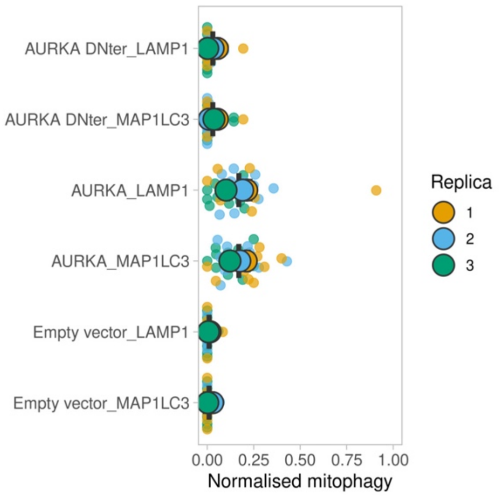


#### Fig. 3A

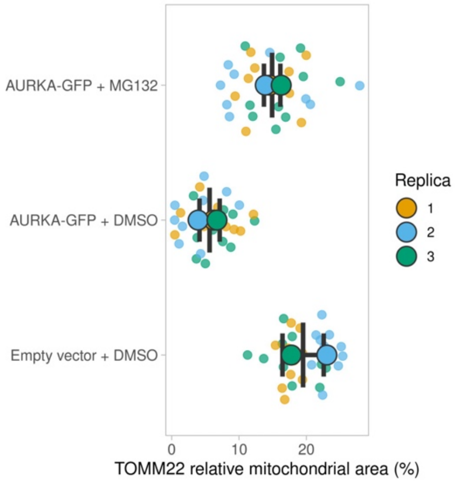


#### Fig. 3C

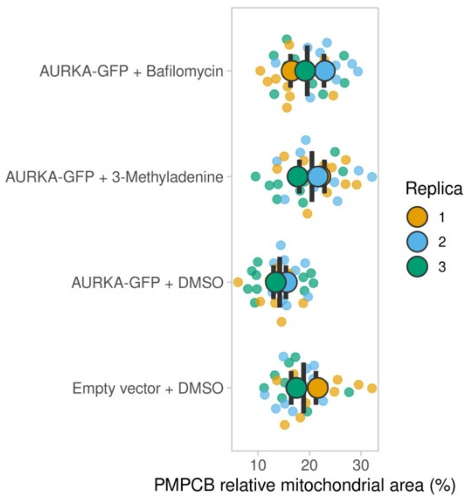


#### Fig. 4A

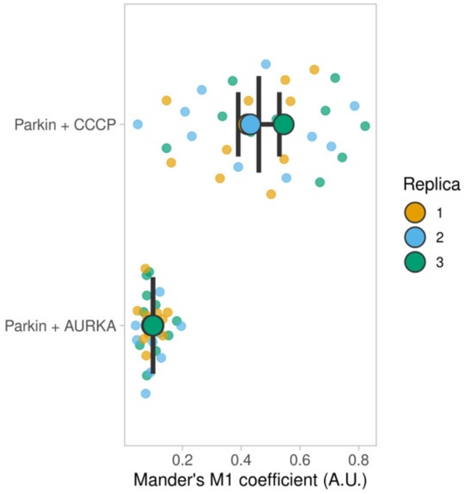


#### Fig. 4B

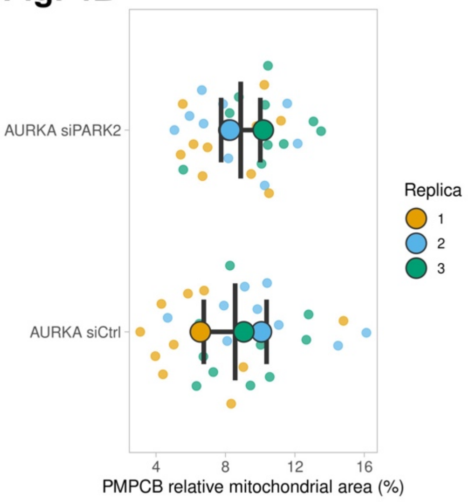


**Supplementary Fig. 8. Representative triplicate data for key quantifications**. *n*=10 individual cells per replicate are shown as pseudocolored dots. Replica means are represented as big, pseudocolored circles, and overall means ± sd are shown for each condition. Graphs were generated with SuperPlotsOfData (https://huygens.science.uva.nl/SuperPlotsOfData/).

#### Fig. 5B

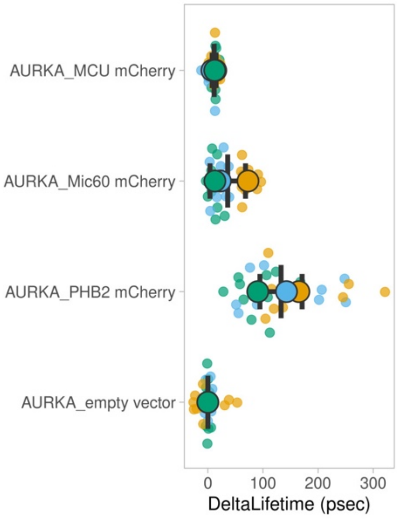


#### Fig.7A

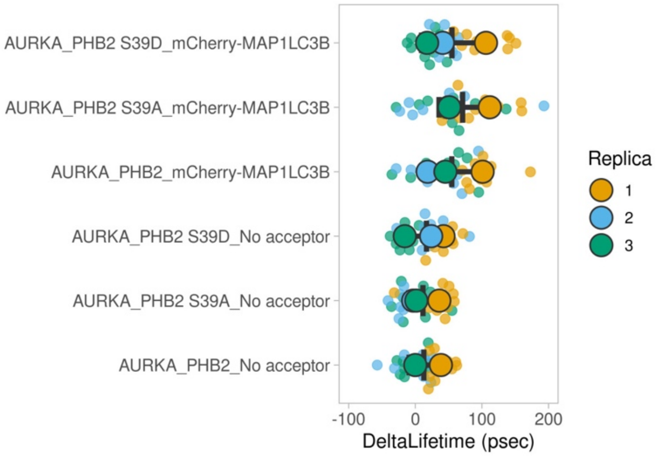


#### Fig. 8C

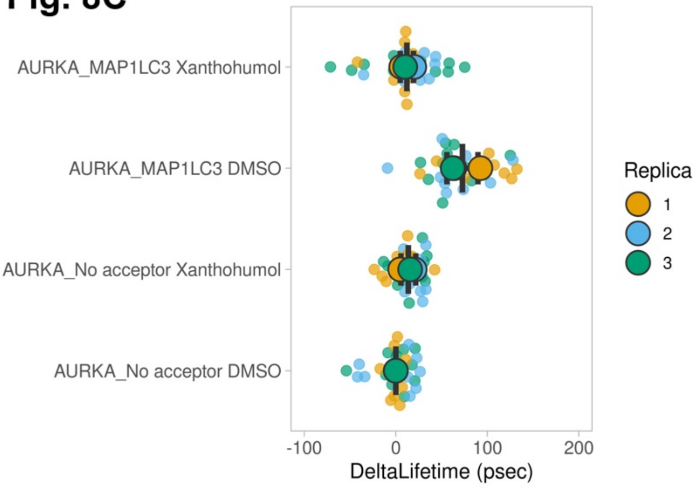


#### Supplmentary Fig. 1A

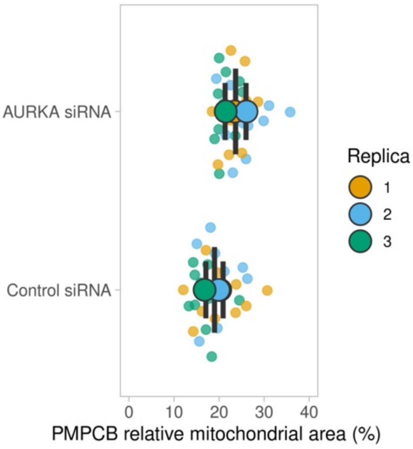


#### Fig. 8D

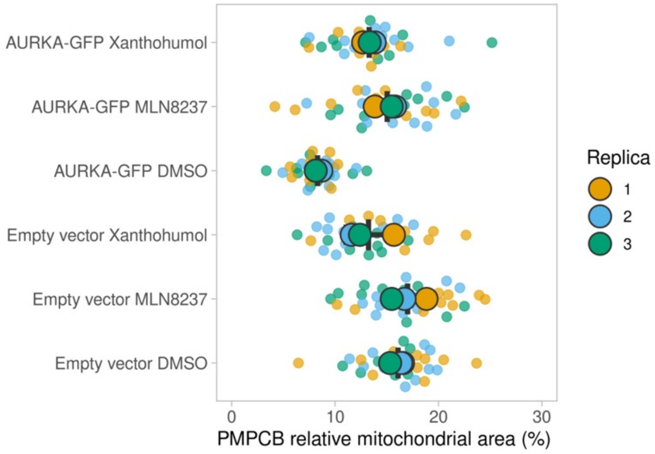


#### Supplmentary Fig. 2

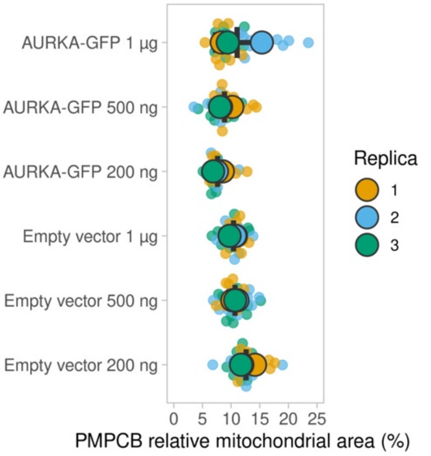


**Supplementary Fig. 9. Representative triplicate data for key quantifications (continued)**. *n*=10 individual cells per replicate are shown as pseudocolored dots. Replica means are represented as big, pseudocolored circles, and overall means ± sd are shown for each condition. Graphs were generated with SuperPlotsOfData (https://huygens.science.uva.nl/SuperPlotsOfData/).

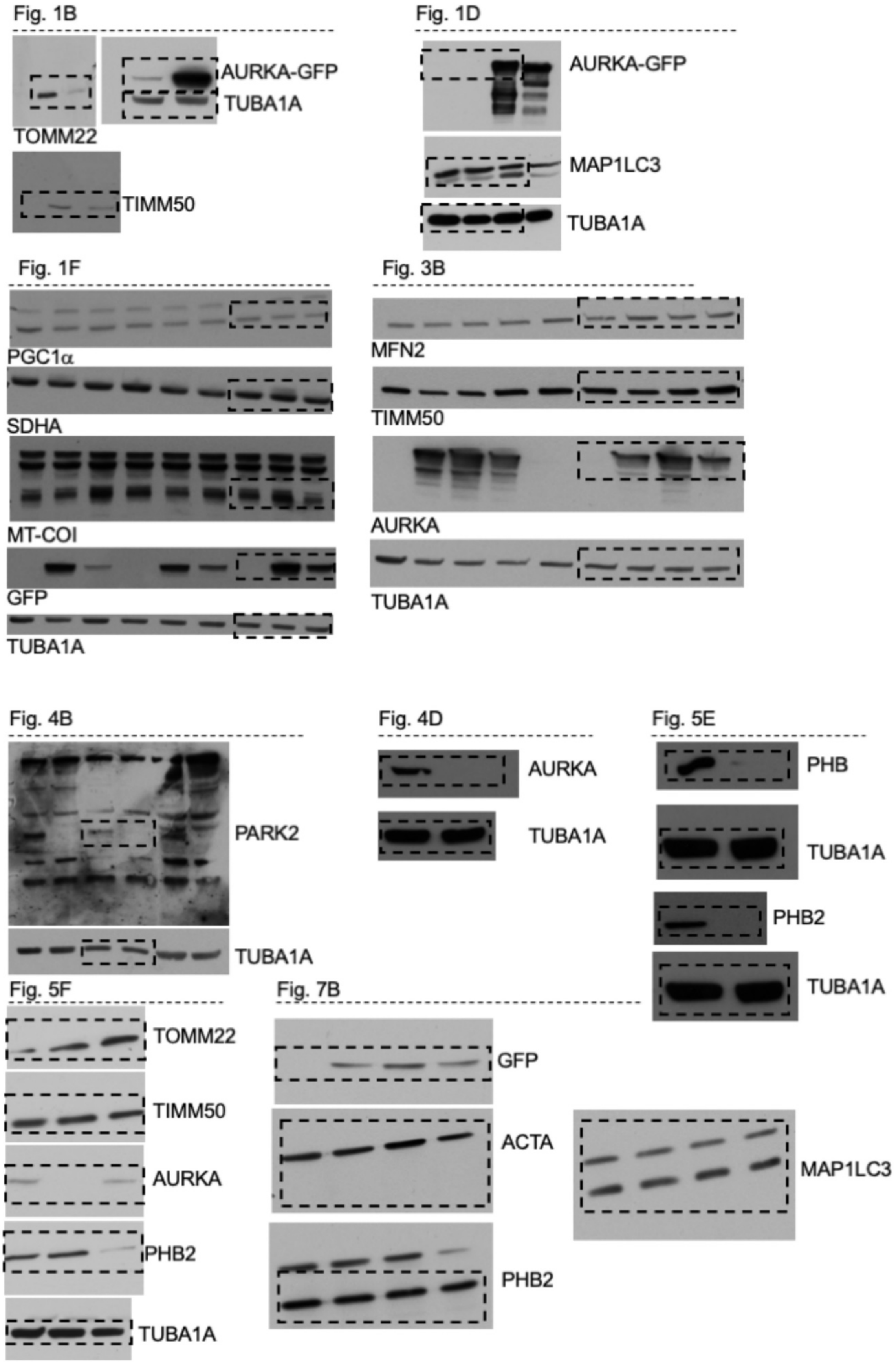


**Supplementary Fig. 10. Representative uncropped blots**. Scans of the full blots used in the study, along with each detected protein and the corresponding figure. The areas enclosed by a dashed rectangle are the portions of the blots integrated in the figure of the study.

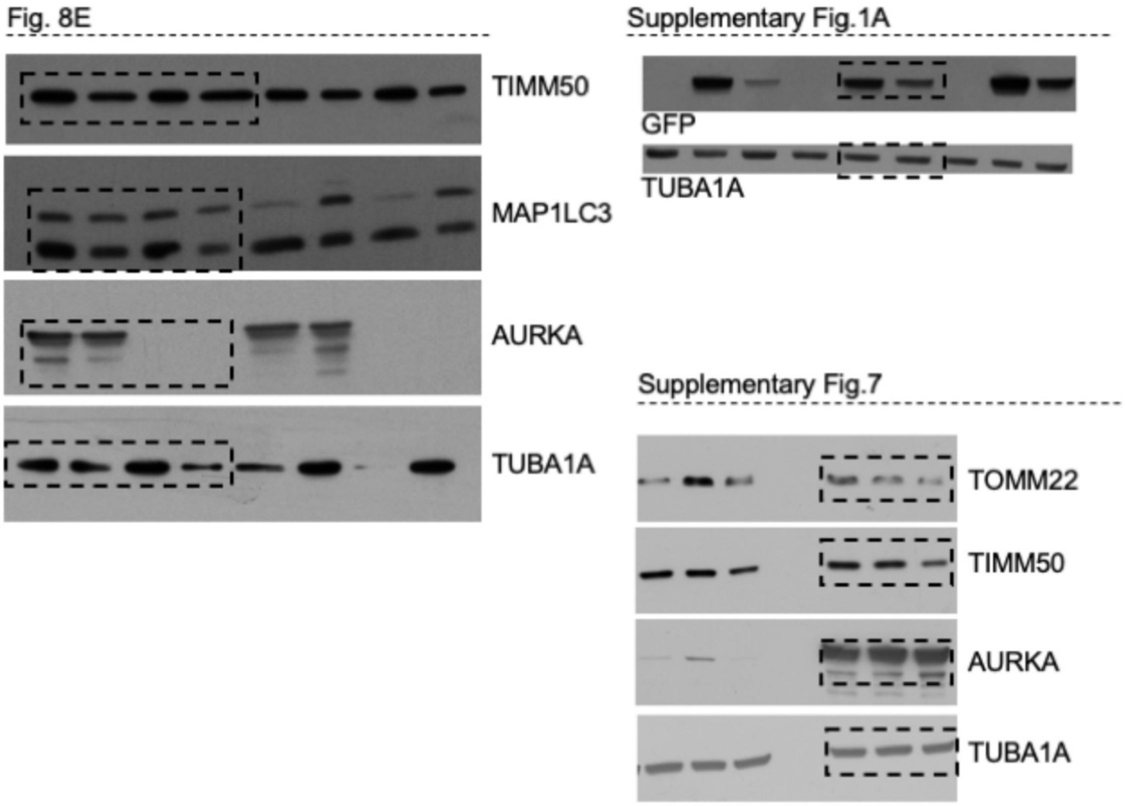


**Supplementary Fig. 11. Representative uncropped blots (continued)**. Scans of the full blots used in the study, along with each detected protein and the corresponding figure. The areas enclosed by a dashed rectangle are the portions of the blots integrated in the figure of the study.

| Plasmid | Cloning sites | Remarks | Primers for mutagenesis (5' to 3'): sense | Primers for mutagenesis (5' to 3'): anti-sense |
|---|---|---|---|---|
| pcDNA 3.1 AURKA 6xHis | | *From Bertolin et al, 2018.* | | |
| pcDNA 3.1 AURKA ΔNter 6xHis | | *the first 30 amino acids of AURKA are lacking; from Bertolin et al, 2018.* | | |
| pcDNA3 AURKA-GFP | | *From Bertolin et al, 2018.* | | |
| pEGFP N1 AURKA ΔNter | | *the first 30 amino acids of AURKA are lacking; from Bertolin et al, 2018.* | | |
| pLKO Puro shControl | | *purchased from Sigma-Aldrich (see materials and methods)* | | |
| pLKO Puro shAURKA | | *purchased from Sigma-Aldrich (see materials and methods)* | | |
| pEGFP N1 mitoGFP | | *From Bertolin et al, 2018.* | | |
| pCMV7 mitomCherry | | *Kind gift of O. Corti, Brain and Spine institute, France.* | | |
| pcDNA 3.1 mCherry-hLC3B | | *Addgene plasmid: #40827* | | |
| pDEST GFP-mCherry MAP1LC3B | | *Kind gift of M. Marinello, Genethon, France* | | |
| pmCherry LAMP1 | | *Addgene plasmid: #45147* | | |
| pEGFP-Parkin WT | | *Addgene plasmid #45875* | | |
| pmCherry MIC60 | XhoI/HindIII | *Kind gift of S. Jakobs, Max Planck Institute, Germany* | | |
| pmCherry MCU | XhoI/HindIII | *Addgene plasmid #31732* | | |
| pmCherry N1 PHB | XhoI/HindIII | *Kind gift of B. Levine* | | |
| pmCherry N1 PHB2 | XhoI/HindIII | | | |
| pmCherry N1 PHB2 S39A | | *obtained from pmCherry N1 PHB2* | cacggtgaacacagcttcgcgcacaccgta | tacggtgtgcgcgaagctgtgttcaccgtg |
| pmCherry N1 PHB2 S39D | | *obtained from pmCherry N1 PHB2* | tccacggtgaacacatcttcgcgcacaccgtagg | cctacggtgtgcgcgaagatgtgttcaccgtgga |
| pcDNA 3.1 (-) PHB2 | NheI/EcoRI | *PHB2 template obtained from pmCherry N1 PHB2* | | |
| pcDNA 3.1 (-) PHB2 S39A | NheI/EcoRI | *PHB2 S39A template obtained from pmCherry N1 PHB2 S39A* | | |
| pcDNA 3.1 (-) PHB2 S39D | NheI/EcoRI | *PHB2 S39D template obtained from pmCherry N1 PHB2 S39D* | | |
| pEGFP N1 PHB2 | AgeI/NotI | *PHB2 template obtained from pmCherry N1 PHB2* | | |
| pEGFP N1 PHB2 S39A | AgeI/NotI | *PHB2 S39A template obtained from pmCherry N1 PHB2 S39A* | | |
| pEGFP N1 PHB2 S39D | AgeI/NotI | *PHB2 S39D template obtained from pmCherry N1 PHB2 S39D* | | |
| pmCherry N1 mitoGFP | XhoI/HindIII | *mitoTandem (mitoGFP fused to mCherry)* | | |

**Supplementary Table 1. List of plasmid vectors used in this study.** Table illustrating the source of each plasmid, cloning sites (when applicable) and primers used for site-directed mutagenesis.

| Antibody | Brand | Catalogue n° | Dilution |
|---|---|---|---|
| Actin | Sigma-Aldrich | A5060 | 1/1000 |
| AURKA clone 5C3 | home made | N.A. (as in Bertolin et al, 2018) | 1/20 |
| GFP | Roche/ Sigma Aldrich | 11814460001 | 1/1000 |
| LC3 | Novus Biologicals | NB100-2331 | 1/1000 |
| MitoFusin2 | Abcam | ab56889 | 1/500 |
| MT-CO1 | Abcam | ab203912 | 1/1000 |
| MT-CO2 | home made | N.A. (From Agier et al, 2012) | 1/2000 |
| PARKIN clone PARK8 | Millipore | MAB5512 | 1/1000 |
| PGC1$\alpha$ | Abcam | ab188102 | 1/1000 |
| PHB | Thermo Scientific | PA5-19556 | 1/500 |
| PHB2/REA | Thermo Scientific | PA5-79817 | 1/1000 |
| SDHA | Abcam | ab137040 | 1/2000 |
| Tim50 | Abcam | ab109436 | 1/2000 |
| Tom22 | Abcam | ab10436 | 1/500 |
| Tubulin alpha clone YL1/2 | Millipore | MAB1864 | 1/5000 |

**Supplementary Table 2. List of primary antibodies used for western blotting procedures.** The table contains the name of the antibody used, the supplier, the catalogue number and the final dilution used

| Genotypes | Name used in publication |
| --- | --- |
| Sca-GAL4, UAS-mito-GFP / UAS-mCherry Atg8a ; + | (*Atg8*; +) |
| Sca-GAL4, UAS-mito-GFP / UAS-mCherry Atg8a ; UAS-*aurA*.Exel / + | (*Atg8*; *aurA* $^{O/E}$) |
| Sca-GAL4, UAS-mito-GFP / UAS-mCherry Atg8a ; *aurA* $^{ST/+}$ | (*Atg8*; *aurA* $^{ST/+}$) |
| Sca-GAL4, UAS-mito-GFP / UAS-mCherry Atg8a ; *aurA* $^{ST/2G}$ | (*Atg8*; *aurA* $^{ST/2G}$) |

**Supplementary Table 3. List of *Drosophila* crossings used in the study.** List of the genotype of the *Drosophila* crossings used in this study, along with the corresponding name used in the publication.
